# BoolDog: integrated Boolean and semi-quantitative network modelling in Python

**DOI:** 10.64898/2026.03.16.711264

**Authors:** Carissa Bleker, Maja Zagorščak, Andrej Blejec, Kristina Gruden, Anže Županič

## Abstract

**Summary:** Boolean and logic-based modeling approaches are well suited for the analysis of complex biological systems, particularly when detailed biochemical and kinetic information is unavailable. In such settings, biological pathways are represented as networks capturing system components and their interactions, providing a simplified yet informative abstraction of system behavior. While the structural topology of these networks is often well characterized, the absence of mechanistic detail limits the applicability of parameter-dependent modeling frameworks. To address this, we present BoolDog, a Python package for the construction, simulation, and analysis of Boolean and semi-quantitative Boolean networks. BoolDog supports synchronous simulation with events, attractor and steady-state identification, network visualization, and the systematic transformation of logic-based models into continuous ordinary differential equation (ODE) systems — enabling the seamless integration of discrete and continuous modeling paradigms. Networks can be imported and exported across standard formats, and BoolDog integrates natively with established Python libraries for network analysis and visualisation, including NetworkX, igraph, and py4Cytoscape. Together, these capabilities provide a flexible, accessible, and interoperable platform for logic-based modeling of complex biological systems.

**Availability and implementation:** BoolDog is implemented in Python and available at https://github.com/NIB-SI/BoolDog/.

## Introduction

Systems biology increasingly relies on computational models to integrate knowledge from heterogeneous sources and to translate qualitative descriptions of cellular processes into executable models. Although detailed kinetic models can provide deep mechanistic insights, they require extensive parameterisation that is often unavailable. For many biological questions, such as exploring perturbations, robustness, or dynamical trends, Boolean and logic-based models are more suitable^1^. These models encode system structure and regulatory logic without requiring detailed reaction kinetics, enabling broad applicability to model gene regulation and signal transduction across disease and developmental processes^2–5^.

In their simplest form, regulatory networks in biology represent entities as nodes and regulatory relationships, such as activation or inhibition, as directed edges. Such networks provide structural insight but lack explicit rules governing state transitions. Boolean networks extend this representation by assigning binary states to nodes (biological entities) and specifying logical update functions defining each node’s state as dependent on its regulators (symbolising the biological function of the relations between nodes). Boolean models are curated in public repositories including Cell Collective^6^, Biomodels^7^, Biodivine^8^, and GINsim^9^. Community efforts such as the CoLoMoTo notebook^10^ have further advanced reproducibility and accessibility by providing a unified Jupyter-based environment integrating a wide range of Boolean modelling tools.

The conversion of regulatory networks into Boolean networks can be non-trivial, owing to ambiguities in combinatorial regulation, interaction strengths, and context-dependent effects. Regulatory networks may also encode multi-component complexes or higher-order interactions that require a hypergraph formalism for correct interpretation. While intuitive and powerful, Boolean models are inherently discrete, yet in reality biological components often exhibit gradations of activity rather than strictly binary behaviour. This motivates continuous simulations of Boolean networks, either stochastically, for example as a continuous-time Markov process as in MaBoSS^11^ or deterministically, by mapping Boolean logic into continuous dynamical ODE systems using smooth activation functions^12^, interpolation schemes^13^, or piecewise-linear differential equations^14^.

Here we present BoolDog, a Python package for the construction, modification, and synchronous analysis of Boolean networks. Additionally BoolDog addresses the challenges above by providing dedicated tools for both regulatory and Boolean models, supporting their interconversion, and implementing established ODE transformation schemes. Together, these capabilities offer a flexible and accessible platform for logic-based modeling of complex biological systems.

## Results

### Tool overview and functionalities

BoolDog is a Python package for the import, modification, visualisation, simulation, and analysis of Boolean networks (Figure 1A). The package accepts Boolean models in Boolnet^15^, SBML-qual^16^, and TabularQual [ref] formats. Regulatory networks encoding activation and inhibition relationships without explicit update logic can be imported from GraphML and SIF files, or directly from NetworkX^17^ and igraph^18^ objects, and are automatically transformed into Boolean models.

**Figure 1:**
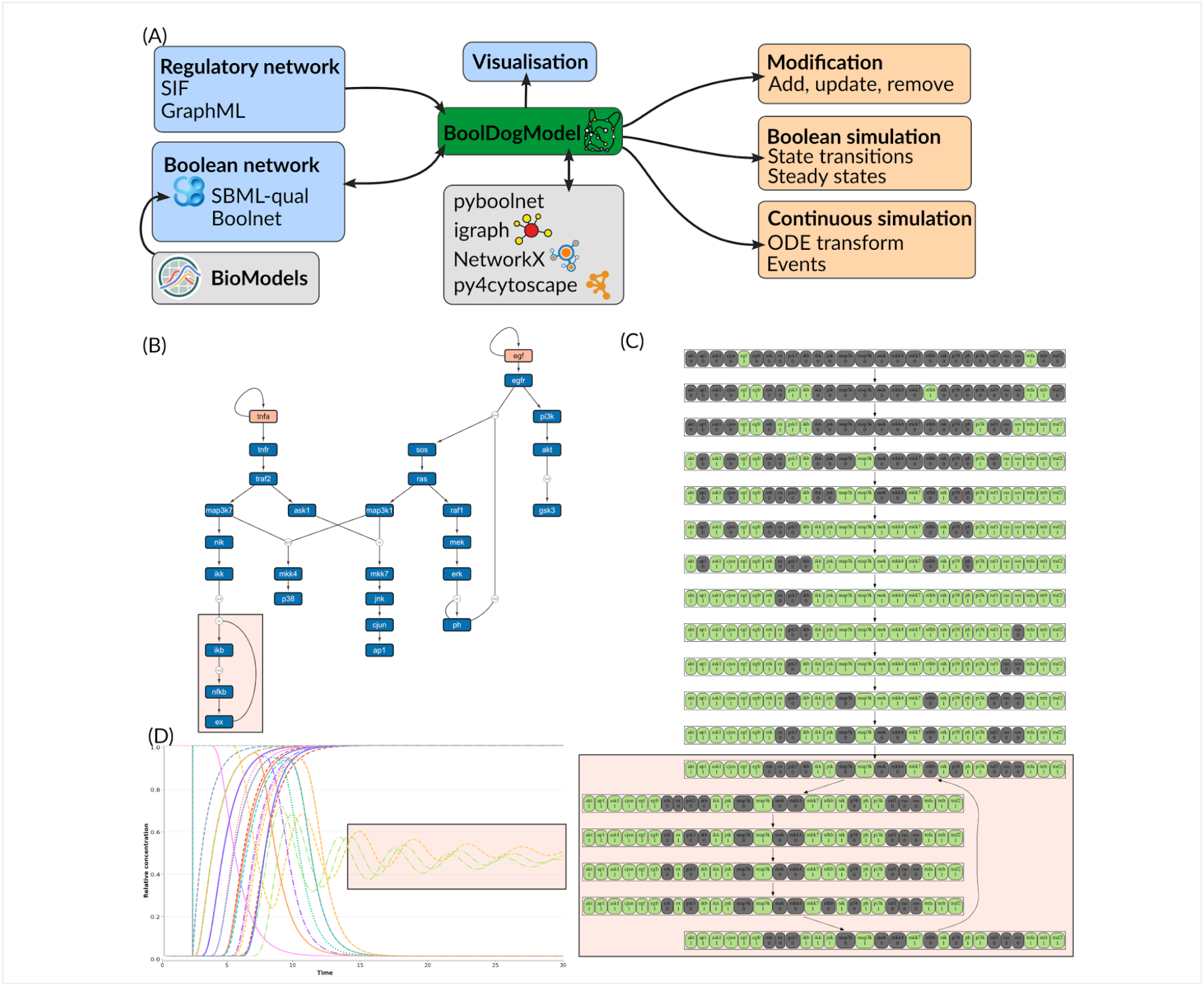
(A) Overview of BoolDog functionalities. (B - D) Case study of EGF and TNFα signalling with (B) Boolean model using the BoolDog - Cytoscape visualisation tool, (C) Boolean simulation (state transition graph) of the model with EGF + TNFα activated (Case ii), showing the attractor landscape, and (D) continuous ODE simulation starting from the inactive steady state, with an event activating EGF and TNFα at pseudo-timepoint 2, showing dampened oscillatory responses. A notebook, showing the process to generate the case study images, as well as an animation of B displaying the transitions in C, is available in the BoolDog documentation: https://nib-si.github.io/BoolDog/gh-pages/tutorials/tutorial-advanced.html

The model can be modified, by adding, removing, or updating nodes, allowing interactive refinement. Visualisation of Boolean models, optionally with a hypergraph representation to represent higher-order interactions, is supported with existing NetworkX or igraph functionalities, or through built in Cytoscape Automation^19^, enabling seamless and interactive visualisation. Models can be exported in BoolNet and SBML-qual formats. Export to additional formats is available through NetworkX and iGraph interoperability.

BoolDog supports synchronous simulation and steady-state and attractor identification to characterise the long-term dynamical behaviour of the network, leveraging PyBoolNet^20^. Continuous dynamics in BoolDog are enabled through two established ODE transformation schemes, SQUAD^21^ and ODEfy^22^, converting Boolean logic into systems of ordinary differential equations and bridging discrete and continuous modeling paradigms. Continuous simulations support events, enabling the modelling of targeted perturbations such as node knockouts or forced activations and inhibitions at defined time points. All major analytical outputs can be visualised directly from BoolDog, supporting both exploratory analysis and the preparation of publication-ready figures. Together, these functionalities provide an integrated workflow from network construction to dynamical analysis and continuous transformation, accessible within the broader Python scientific ecosystem.

### Comparison with existing tools

A number of tools support Boolean network modelling, including GINsim^9^, PyBoolNet^20^, BoolNet^15^, and MaBoSS^11^, among others; a recent comprehensive comparison of is provided by Saalfeld et al.^23^ In benchmarking approaches for gene regulatory networks, tools have also been developed to generate continuous data from gene regulatory networks, based on ODE models of transcriptional regulation^24–26^. These tools however only consider the specific case of generating synthetic gene expression data, and are not generalised to signalling networks as a whole. Here we focus on tools that, like BoolDog, support semi-quantitative ODE-based transformations of generic Boolean networks: ODEfy^22^, SQUAD^27^, Genetic Network Analyzer (GNA)^14^, and JimenaE^28^. These tools present limitations in terms of accessibility, maintenance, and interoperability (Table 1). ODEfy is implemented in MATLAB with a non-commercial license and has not been actively maintained. SQUAD is a Java GUI application distributed as a binary-only package (compiled against Java 1.6) under a non-commercial license with no source code available, making deployment and reproducibility challenging. GNA takes a related but distinct approach, modelling regulatory networks as piecewise-linear differential equation systems defined by qualitative inequality constraints; while feature-rich, including attractor search and temporal logic verification, it operates in its own modelling formalism rather than on Boolean networks directly, and is distributed as a Java GUI application with no Python interoperability and no source code available. JimenaE (and its predecessors Jimena^29^, Jimena2^30^, JimenaC^31^), while the most feature-rich of the three, is also Java-based and requires MATLAB for ODE routines. None of the three tools offer Python-native workflows, integration with modern network analysis libraries, access to curated model repositories, or persistent identifiers for FAIR-compliant software distribution.

**Table 1:** Comparison of BoolDog with existing semi-quantitative Boolean modelling tools.

|  |  | BoolDog | ODEfy | SQUAD | GNA | Jimena* |
| --- | --- | --- | --- | --- | --- | --- |
| Functionality | Regulatory network import | ✓ | ✓ | ✓ | — | ✓ |
|  | Boolean network import | ✓ | ✓ | — | — | ✓ |
|  | Model visualisation | ✓ | — | ✓ | ✓ | ✓ |
|  | Network-to-Boolean conversion | ✓ | ✓ | ✓ | — | ✓ |
|  | Boolean simulation | ✓ | — | ✓ | — | ✓ |
|  | Boolean attractor/steady-state analysis | ✓ | ✓ | ✓ | — | ✓ |
|  | ODE transformation | SQUAD, ODEfy | ODEfy | SQUAD | piece-wise linear | SQUAD, ODEfy |
|  | Event-based ODE simulation | ✓ | — | ✓ | — | ✓ |
|  | ODE attractor/steady-state analysis | — | ✓ | ✓ | ✓ | ✓ |
|  | SBML-qual import/export | ✓ | — | — | ✓ | — |
|  | BioModels access | ✓ | — | — | — | — |
| Usability | Python-native | ✓ | — | — | — | — |
|  | Last release | 03/2026 (PyPi) | 10/2019 (GitHub) | 11/2007 (Suppl) | 07/2015 (web) | 07/2022 (Suppl) |
|  | Cytoscape integration | ✓ | — | — | — | — |
|  | NetworkX/igraph integration | ✓ | — | — | — | — |
|  | GUI | — | partial‡ | ✓ | ✓ | ✓ |
|  | Documentation | ✓ | ✓ | partial | ✓ | partial |
|  | Tutorials or examples | ✓ | ✓ | ✓ | ✓ | ✓ |
|  | Open license | GPL-3.0 | ✗ non-commercial | ✗ non-commercial | ✗ non-commercial | LGPL-3.0 |
|  | Source code available | ✓ | ✓ | — | — | — |
|  | Persistent identifier | ✓ | — | — | — | — |

\*Jimena Software considers Jimena, Jimena2, JimenaC, JimenaE; ‡ODEfy includes a MATLAB GUI

BoolDog addresses these gaps by providing a fully Python-native, actively maintained, and openly licensed platform that integrates Boolean simulation, semi-quantitative ODE transformation via both SQUAD and ODEfy schemes, and event-based continuous simulation within a single package. Its native interoperability with NetworkX, igraph, and Cytoscape, together with direct BioModels access and Zenodo-archived versioned releases, distinguishes it from existing tools in both functionality and usability. The one area where BoolDog does not yet match the capabilities of ODEfy and JimenaE is ODE-domain attractor and steady-state analysis, which remains an area for development.

### Case study: EGF and TNFα signalling

To demonstrate BoolDog’s core functionality, we applied it to a published Boolean model of EGF and TNFα mediated signalling^16^. BoolDog was used to retrieve the model directly from the BioModels repository (BIOMD0000000562) and imported in SBML-qual format. The model comprises 28 nodes representing key signalling components including EGFR, TNFR, NF-κB, ERK, JNK, and p38. BoolDog was used to interactively visualise the model in Cytoscape as a logic circuit, explicitly encoding the Boolean update rules for each node (Figure 1B).

Synchronous Boolean simulations were performed from two initial conditions: a fully inactive state, and a state in which both EGF and TNFα inputs were activated (conforming to cases (i) and (ii) in Chaouiya et al. ^16^). In the inactive case, the network converges to a single inactive steady state, whereas dual stimulation with EGF and TNFα leads to a distinct attractor landscape reflecting downstream pathway activation (Figure 1C), consistent with the dynamics reported in the original publication.

The Boolean model was subsequently transformed into a continuous ODE system using the normalised HillCube scheme. Starting from the inactive steady state identified by the above Boolean analysis, an event activating both EGF and TNFα inputs was introduced mid-simulation. The resulting trajectories reveal dampened oscillatory responses in downstream signalling components (Figure XD), capturing transient dynamics that are not accessible in the discrete representation. The transition from discrete to continuous dynamics was achieved with a single function call, illustrating the accessibility of BoolDog’s hybrid modeling workflow. These results illustrate BoolDog’s integrated workflow from model import and visualisation through Boolean analysis and continuous semi-quantitative simulation.

### Conforming to FAIR principles

BoolDog is developed in accordance with the FAIR4RS principles^32^. The package is assigned a persistent identifier via PyPI and versioned releases are archived on Zenodo^33^, each receiving a distinct DOI (F1, F1.2), with metadata and documentation publicly available and indexable (F2–F4). The source code and all associated metadata are openly retrievable via standard protocols (A1, A1.1), and Zenodo archiving ensures long-term accessibility even if the primary repository becomes unavailable (A2). BoolDog reads and writes established community formats including SBML-qual and integrates natively with widely used Python libraries for network analysis (I1, I2). The software is released under a GPL-3.0 license, accompanied by provenance, versioned releases, tutorials, and API documentation (R1–R3).

## Discussion

BoolDog fills an important gap in the logic-based modelling landscape by providing a single interface for the construction, Boolean analysis, continuous transformation and event-based continuous simulation of Boolean and regulatory networks, advancing the ability to model complex biological systems. BoolDog’s Python-native architecture means that missing functionality can be readily addressed by leveraging the extensive Python scientific ecosystem, or implemented directly by users familiar with Python, without requiring access to proprietary environments or compiled binaries. As an openly developed package, it is also a candidate for future inclusion in the Colomoto notebook^10^, further contributing to reproducible and accessible Boolean modelling workflows.

## Methods

### Implementation

BoolDog is implemented in Python (≥3.12) and is available under a GPL-3.0 license via PyPI and GitHub (https://github.com/NIB-SI/BoolDog), with versioned releases archived on Zenodo^33^. It can be installed using pip (`pip install booldog`). Full documentation, including installation instructions, API reference, and tutorials, is available at https://nib-si.github.io/BoolDog/.

BoolDog network objects are interoperable with igraph^18^ and NetworkX^17^. SBML-qual^16^ and TabularQual [ref] interoperability are supported by python-libsbml^34^ and the TabularQual converter [ref] respectively. Boolean simulations and attractor and steady-state analysis utilise Answer Set Programming (ASP) as implemented in PyBoolNet^20^. Continuous ODE simulations are implemented using SciPy^35^ and NumPy^36^. Visualisations utilise matplotlib^37^ and pygraphviz^38^, and network visualisation in Cytoscape^39^ is automated using the py4cytoscape library^40^. Interoperability and visualisation dependencies are available as optional installation extras to minimise the core installation footprint.

## Data availability

No data was generated in this study.

## Code availability

BoolDog source code is available at https://github.com/NIB-SI/BoolDog. All analyses reported here can be reproduced using notebooks available in the GitHub repository.

## Acknowledgements

BoolDog development with funding and support from the European Union’s Horizon 2020 research and innovation programme under grant agreement No. 862858 ([ADAPT](https://adapt.univie.ac.at/)), the [Slovenian Research and Innovation Agency](https://www.arrs.si/en/) (ARIS) under grant agreements No. P4-0463, No. GC-0001, No. J4-1777, No. Z4-50146, as well as supporting research infrastructure used in this work through grant IO-0004. We also acknowledge the Slovenian node of ELIXIR (ELIXIR-SI) for providing access to computational infrastructure.

## Author contributions

**CB**: Methodology, Software, Validation, Visualization, Funding acquisition, Writing - Original Draft. **MZ**: Methodology, Software. **AB**: Methodology, Software. **KG**: Funding acquisition, Conceptualisation Supervision. **AŽ**: Conceptualization, Supervision, Funding acquisition. All authors: Writing - Review & Editing.

## Supplementary information

N/A

